# The road to sorghum domestication: evidence from nucleotide diversity and gene expression patterns

**DOI:** 10.1101/2020.11.17.386268

**Authors:** Concetta Burgarella, Angélique Berger, Sylvain Glémin, Jacques David, Nancy Terrier, Monique Deu, David Pot

## Abstract

Native African cereals (sorghum, millets) ensure food security to millions of low-income people from low fertility and drought-prone regions of Africa and Asia. In spite of their agronomic importance, the genetic bases of their phenotype and adaptations are still not well understood. Here we focus on *Sorghum bicolor*, which is the fifth cereal worldwide for grain production and constitutes the staple food for around 500 million people. We leverage transcriptomic resources to address the adaptive consequences of the domestication process. Gene expression and nucleotide variability were analyzed in 11 domesticated and 9 wild accessions. We documented a downregulation of expression and a reduction of diversity both in nucleotide polymorphism (30%) and gene expression levels (18%) in domesticated sorghum. These findings at the genome-wide level support the occurrence of a genetic bottleneck in the domestication history of sorghum, although several genes showed also patterns consistent with the action of selection. Nine hundred and forty-nine genes were significantly differentially expressed between wild and domesticated gene pools. Their functional annotation points to metabolic pathways most likely contributing to the sorghum domestication syndrome, such as photosynthesis and auxin metabolism. Coexpression network analyses revealed 21 clusters of genes sharing similar expression patterns. Four clusters (totalizing 2449 genes) were significantly enriched in differentially expressed genes between the wild and domesticated pools and two were also enriched in domestication and improvement genes previously identified in sorghum. These findings reinforce the evidence that domestication and improvement do not only affect the behaviors of a few genes but led to a large rewiring of the transcriptome during the domestication event and the improvement process. Overall, these analyses pave the way towards the identification of key domestication genes valuable for genetic resources characterization and breeding purposes.

## 1. Introduction

Domestication is an evolutionary process in which species evolve dramatic functional and phenotypic changes under human selection for characters that fit the agricultural environment and human necessities (such as taste, yield, cultivation, harvesting and storage practices). The resulting domesticated forms are different from unmanaged wild populations and generally unable to reproduce and survive in the wild. The suite of traits that have been modified in the domesticated forms are collectively referred to as “domestication syndrome” (Allaby, 2014). In plants they typically include characters related to seed dispersal (e.g. reduction of dehiscence and dormancy), plant architecture (e.g. branching) and properties of the harvested part (such as size, shape, nutritional content of seed, fruit and tubers). Around 250 plants are estimated to have experienced domestication (Dirzo and Raven, 2003), with grasses (family Poaceae) having contributed at least 48 species (Glémin and Bataillon, 2009).

The extensive phenotypic changes induced in domesticated species are associated with genetic changes, which can be put in evidence by comparing domesticated and wild forms. According to domestication models, an important genome wide signature of the domestication process is a genetic bottleneck experienced by the domesticated forms, which would be due to a small sampling from the wild species associated with the selection for favorable traits (Doebley et al., 2006; Meyer and Purugganan, 2013). This general reduction of genetic diversity is expected to be stronger for annual than perennial species and in autogamous than allogamous crops (Gaut et al., 2015). Experimental evidence supports this hypothesis, as a reduction of genetic diversity has been found in major domesticated species when compared with their wild progenitors, e.g. maize (Hufford et al., 2013), rice (Caicedo et al., 2007; Huang et al., 2012; Nabholz et al., 2014), soybean (Lam et al., 2010) and tomato (Koenig et al., 2013; Sauvage et al., 2017).

The domesticated genome is expected to show also other genome-wide signatures when compared with the wild progenitor, such as increased linkage disequilibrium (e.g. Wright et al., 2005; Zhou et al., 2015) and increased mutation load, i.e. a genome-wide accumulation of weakly deleterious genetic variants that reduces the fitness of the domesticated forms compared to the wild counterpart. This latter phenomenon has been called ‘domestication cost’ (Lu et al., 2006) and has been observed in several domesticated animal and plant species (Moyers et al., 2018), including rice (Nabholz et al., 2014; Liu et al., 2017), maize (Wang et al., 2017) and sunflower (Renaut and Rieseberg, 2015).

Besides genome-wide effects, the specific genomic regions involved in the phenotypic traits that distinguish domesticated from wild forms are expected to show stronger molecular genetic signatures than the rest of the genome. Although the genetic mechanisms under many complex domestication traits are still unknown, several major domestication genes have been identified along with their causative variation, which includes single nucleotide mutations, copy number polymorphism or indels changes (Kantar et al., 2017; Purugganan, 2019). These genes include both coding and regulatory regions, indicating that changes in expression levels may be as important as protein modifications (Meyer and Purugganan 2013, Olsen and Wendel 2015). Supporting this hypothesis, recent surveys of gene expression patterns have put in evidence significant differences between the domesticated and the wild pools (e.g. Bellucci et al., 2014; Page et al., 2019). For example, in maize and tomato the rewiring of gene expression levels has been documented for some functional networks (Swanson-Wagner et al., 2012; Sauvage et al., 2017) up to shutdown complete pathways (Itkin et al., 2013), suggesting that the expression changes induced by domestication may have targeted metabolic pathways more than major effect genes (Sauvage et al., 2017). Domestication has also involved loss of connectivity within co-expression networks for some genes (Swanson-Wagner et al., 2012).

We focus here on sorghum (*Sorghum bicolor* (L.) Moench), the world’s 5^th^ cereal for grain production. Being staple food for more than 500 million people, sorghum plays a key role in food security in Africa and Asia. Aside from providing food for human consumption, sorghum is widely cultivated in all continents as a source for feed, fiber and energy (Paterson et al., 2009). In the current context of climate change and resource depletion, sorghum is also a cereal with great potential for future agriculture, because it outperforms other crops under low-input and stressful conditions (Hasan et al., 2017).

Sorghum was first domesticated in North-East Africa, likely around 6000 years ago (Winchell et al., 2017, 2018). The domesticated phenotype is distinguishable from the wild forms in many morphological, physiological and phenological traits, including non-shattering grains, higher seed size, different panicle and plant architecture. Some genes and gene families have been identified as likely targets of the domestication and/or the improvement processes. Notably, the loss of seed shattering has been related to mutations in the *shattering1* (*sh1*) locus, which has likely undergone parallel selection in sorghum, rice and maize (Lin et al., 2012). Selection scans and phenotype-genotype association studies have identified other candidate genes. These include known cereal domestication genes Tb1 (*teosinte branched1*) and ba1 (*barren stalk1*), involved in plant architecture (Mace et al., 2013; Lai et al., 2018), as well as genes associated with grain features (e.g. Psy1, *phytoene synthase1*), photoperiod sensitivity (early maturity Ma genes) and plant height (Dw genes) (Fernandez et al., 2008; Mace and Jordan, 2010; Mace et al., 2013). Genes within key biosynthesis pathways have also been highlighted, tagging in particular the metabolisms of starch (ss1 and sbe3 genes, Campbell et al., 2016), grain size and weight (Tao et al., 2017, 2018).

In spite of its importance, there is still a limited knowledge about the genetic determinants of sorghum traits and adaptive potential. Comparing the domesticated form with its wild progenitor is particularly informative to understand the changes that accompanied the transition from the wild to the crop. By taking the wild pool as reference, we can quantify the genetic changes we observe in the domesticate both at the genome-wide scale and at the individual gene scale. Despite the simplicity of this approach, just a handful of studies have included wild accessions in analyses of sorghum crop diversity and evolution in a comparative manner. These works have analysed variation at allozyme markers (Aldrich et al., 1992), RFLP loci in the nuclear and chloroplast genomes (Aldrich and Doebley, 1992), nuclear microsatellite markers (Casa et al., 2005; Billot et al., 2013) and nucleotide sequence (Hamblin et al., 2005; Mace et al., 2013). They have all reported a global loss of genetic diversity and for some of them evidence of directional selection in the genome of landraces and improved germplasm.

In this work, we address the genetic bases of the transition from wild to domesticate in sorghum by analysing the transcriptome expressed in leaves, flowers and maturing seeds in wild and domesticated accessions. Compared to nucleotide diversity screening, transcriptome analysis provides twofold information, about changes in the nucleotide sequence of coding genes and changes in their levels of gene expression. Here we perform complementary analyses with three main objectives: i) to quantify the effect of domestication on nucleotide polymorphism, by looking at gene diversity and differentiation patterns; ii) to unveil the effect of domestication on patterns of gene expression and document changes in single genes and gene networks; iii) to assess the congruence and divergence of polymorphism and expression patterns to identify genes and pathways under selection and responsible for the evolution of the domesticated phenotype.

## 2. Materials and Methods

### 2.1 Plant material

A data-set of 20 sorghum accessions were examined in this study. Accessions were selected to represent the diversity of the domesticated (n=11) and the wild relative (n=9) gene pools. Details on the accessions are provided in Supplementary Table S1, as well as the correspondence of accession names with published works analysing accessions from this data-set (Clément et al., 2017; Ranwez et al., 2017). All the genotypes were grown in the greenhouse until the mature grain stage. Tissue samples were harvested from three organs: mature leaves (fourth rank below the flag leaf), inflorescence (at the anthesis stage) and maturing grains (in average 25 days after anthesis).

### 2.2 RNA preparation and sequencing

Each tissue was considered independently for RNA extraction, and RNA were then pooled for each accession prior to sequencing. RNA extraction, Illumina library preparation and sequencing conditions are detailed in Sarah *et al*. (2017). A mixture of 65% RNA from the inflorescence, 15% from leaves and 20% from grains for each accession was sequenced using Illumina mRNA-seq, paired-end protocol on a HiSeq2000 sequencer (one run for each genetic pool). The paired-end reads, in the illumina FASTQ format, were cleaned using cutAdapt (Martin, 2011) to trim read ends of poor quality (q score below 20) and to keep only those with an average quality above 30 and a minimum length of 35 base pairs (Sarah et al., 2017). Those data are freely available on the NCBI Sequence Read Archive database (SRA codes listed in Table S1).

### 2.3 Polymorphism analysis

Reads were mapped on sorghum genome version 1.4 with the BWA software (Li and Durbin, 2009), allowing at most three mismatches between a given read and the reference by Sarah *et al*. (2017). To perform genotype calling, we followed the procedure Nabholz *et al*. (2014). We excluded reads with more than two insertions/deletions (indels) or with indels larger than 5 bp or mapping on different transcripts. ORFextractor (available at http://fr.softwaresea.com/download-ORF-Extractor-10449769.htm) was used to extract coding sequences for further processing. The read2snp software (Gayral et al., 2013) was used to call genotypes and filter out paralogs. Following the approach of Nabholz *et al*. (2014) for non-outcrossing species, we did a first run setting the fixation index *F* = 0, then estimated *F* from the data and rerun the program once with the new estimated value. We note that the fixation index calculated here accounts for the mating system and potential substructure in each sample. For each individual and each position, we only kept genotypes with a minimum coverage of 10x and with posterior probability higher than 0.95. Otherwise, data was considered missing. The contigs obtained with reads2snp were further filtered for the downstream analysis.

The structure of genetic diversity revealed through the genotype calling steps was first explored using principal component analysis (PCA) with the function pca in package LEA v. 3.8 (Frichot and François, 2015). For this, we filtered the 24474 contigs common to the wild and domesticated pools by discarding positions called in less than 5 individuals for each genetic pool, ending up with 11699 contigs and 54594 polymorphic loci. The same dataset was analyzed with popGenome (Pfeifer et al., 2014) to calculate population genomic statistics of diversity and differentiation: nucleotide diversity *π* and Tajima’s *D* per pool, *F*_ST_ between pools. The fixation index *F* and polymorphism at synonymous (*π*s) and nonsynonymous (*π*n) sites were calculated with dNdSpiNpiS (available at https://kimura.univ-montp2.fr/PopPhyl/. In dNdSpiNpiS analysis, we started from 26552 and 26274 contigs in domesticated and wild pools respectively, and discarded positions called in less than 5 individuals for each genetic pool, ending up with 13231 and 11397 contigs (Table 1).

**Table 1.** Characteristics of the data sets, coding sequence polymorphism and differentiation in wild and domesticated accessions of *Sorghum bicolor*.

|  | Wild | Domesticate |
| --- | --- | --- |
| **No. of accessions** | 9 | 11 |
| **No. of contigs retained by reads2snp** | 26274 | 26552 |
| **No. of contigs analysed with dNdSpiNpiS** | 11397 | 13231 |
| **Mean length of retained contigs<sup>a</sup>**<br>**[95% quantiles]** | 1610<br>[326; 3882] | 1596<br>[312; 3830] |
| **No. of polymorphic sites<sup>a</sup>** | 46147 | 41179 |
| **F [95%CI]<sup>a</sup>** | 0.42<br>[0.408; 0.424] | 0.99<br>[0.987; 0.989] |
| **mean $\pi_S$<sup>a</sup>**<br>**[95% CI]** | 4.1<br>[3.97; 4.17] | 3.3***<br>[3.15; 3.35] |
| **mean $\pi_N$<sup>a</sup>**<br>**[95% CI]** | 0.7<br>[0.64; 0.67] | 0.5***<br>[0.48; 0.51] |
| **mean $\pi_N/\pi_S$<sup>a</sup>**<br>**[95% CI]** | 0.16<br>[0.156; 0.166] | 0.15<br>[0.147; 0.158] |
| **No. of contigs analysed with PopGenome** | 11699 |  |
| **Length of retained contigs<sup>b</sup>**<br>**Mean [95% quantiles]** | 743 [0; 2600] |  |
| **No. of polymorphic sites<sup>b</sup>** | 32712 | 24453 |
| **Tajima's D<sup>b</sup>**<br>**Median [95% quantiles]** | -0.4<br>[-1.6; 1.6] | -0.39***<br>[-1.8; 1.8] |
| **mean $\pi$ [95% CI]<sup>b</sup>** | 1.6<br>[0; 6.7] | 1.1***<br>[0; 5.4] |
| **Mean $\pi_{CROP}/\pi_{WILD}$ [95% quantiles]** | 0.65 [0; Inf] |  |
| **Mean $F_{ST}$ [95% quantiles]<sup>b</sup>** | 0.07 [0; 0.42] |  |

F: fixation index (Weir & Cockerham 1984).
$\pi$ : nucleotide diversity ( $\times 10^{-3}$ ). All polymorphic positions ( $\pi$ ), synonymous sites ( $\pi_S$ ), non-synonymous sites ( $\pi_N$ )
$F_{ST}$ : pairwise differentiation (Hudson, R. R., M. Slatkin, and W.P. Maddison (1992))
<sup>a</sup> calculated with dNdSpiNpiS,
<sup>b</sup> calculated with popGenome
\*\*\* Kolmogorov-Smirnov test for difference between domesticated and wild sorghum distributions, two-side p-value < 2.2e-16

### 2.4 Differential expression analysis

Transcript expression levels were estimated with the new-Tuxedo pipeline (Pertea et al., 2016). Firstly, for each accession, RNA-seq reads were mapped on the sorghum genome assembly Sbicolor_313_v3.1 using Hisat2 (Kim et al., 2015). Genes and transcripts were assembled and quantified with stringtie (Pertea et al., 2015), using the reference annotation file to guide the assembly process. The output includes expressed reference transcripts as well as any novel transcripts that were assembled. Gffcompare (https://github.com/gpertea/gffcompare) was used to compare transcripts with the reference annotation (gene and transcript predictions) and to identify new genes/transcripts. Assembling and enriched annotation files were used to estimate abundance with stringtie (script provided as Supplementary Material).

The identification of differentially expressed genes between the wild and the domesticated pools was done with edgeR (Robinson et al., 2010) under the R environment (R Core Team, 2018). Only genes assayed in at least 5 accessions and with at least 1 CPM (count per million) in each accession were included in subsequent analyses, considering that the expression level could not be reliably estimated for genes that did not meet these criteria. Since the minimum library size was 12M, one CPM corresponded to 12 reads. A total of 24646 genes passed these filters. To account for gene-specific expression biases at the individual level, a normalization step (trimmed mean of M-values (TMM) between each pair of samples) was performed (Robinson and Oshlack, 2010). The structure of the sample based on the gene expression levels was explored with a PCA on the normalized counts with the function dudi.pca of the R package ade4 (Dray and Dufour, 2007). Diversity in expression within each of domesticated and wild groups was quantified with the coefficient of variation (CV) in read counts. Domesticate-wild differences in CV were assessed in different gene categories (candidate and non-candidate genes under selection), to assess if the CV loss observed in domesticated sorghum was due to selection in the domesticated pool or drift associated to the domestication/improvement process, following the approach of Bellucci *et al*. (2014). Candidate genes were identified by extreme values of the polymorphism and differentiation statistics, non-candidates were all the other genes. The significance of the difference was assessed with 1000 random resamplings.

To identify differentially expressed (DE) genes, gene-wise exact tests for differences in the means between the two groups (wild and domesticated sorghum) were performed. A False Discovery Rate approach was applied to correct for multiple tests. Thresholds of 0.05 and 0.01 FDR were considered to declare DE genes. At 0.05 FDR, we declared a number of DE genes twofold higher than at 0.01 FDR, but the ratio of up- and downregulated genes was qualitatively similar, as well as the main categories of Gene Ontology annotation (see below). We therefore only report the results for the stringent FDR of 0.01.

### 2.5 Polymorphism/expression comparisons

We compared patterns of gene expression and polymorphism to identify genetic signatures potentially linked to selection and the process of domestication. Since expression level can affect the intensity of selection (Drummond et al., 2005; Park et al., 2013; Nabholz et al., 2014), we looked at the correlation between polymorphism (synonymous, *π*_S_, non-synonymous, *π*_N_ and the ratio *π*_N_/*π*_S_) and expression in each gene pool, and tested for species difference by performing an analysis of variance including pools, expression level, and their interaction. Gene expression was estimated as the mean of normalized read counts across accessions within each gene pool. For these analyses, polymorphism values and expression level were log transformed.

We also compared the levels of diversity and differentiation between DE and non-DE genes, to detect signatures exclusive of DE genes and consistent with directional selection in the wild or the domesticated pool. In particular, we tested if DE genes were enriched for lower polymorphism values, or extreme values of Tajima’s D or *F*_ST_. Domestication could have driven selection only on expression, with no changes in the nucleotide sequence. Thus, we also tested if expression divergence of DE genes was stronger than expected given their genetic divergence by testing if DE genes were enriched for *F*_ST_. = 0.

### 2.6 Enrichment of different functional categories

We examined the functional characterization of DE genes and assessed if particular gene categories or functions were overrepresented in DE vs non-DE genes, which could be the result of selection. The categories assessed included the Gene Ontology (GO) categories, Sorghum transcription factors (retrieved from Plant Transcription Factor Database http://planttfdb.cbi.pku.edu.cn/index.php?sp=Sbi), stably expressed genes in sorghum (Shakoor et al., 2014), different metabolic pathways (Rhodes et al., 2014, 2017; Shakoor et al., 2014), domestication and improvement genes from published studies (Mace et al., 2013; Lai et al., 2018). Enrichment in the different functional categories was tested with Fisher’s exact test on 2×2 contingency tables. GO annotation per gene was obtained from Sobic 3.1 genome annotation. The correspondence among gene models for the different versions of the Sorghum genome were retrieved with the conversion tool of the Grass Genome Hub (https://grass-genome-hub.southgreen.fr/pseudomolecules-converter). Functional annotation of novel assembled genes was done using Blast2GO (Götz et al., 2008). GO enrichment analysis was performed with topGO (Alexa et al., 2006) using the “classic” algorithm and Weighted Fisher’s test to generate p-values. Map plots of non redundant categories were done with REVIGO (Supek et al., 2011).

### 2.7 Gene expression networks

Gene co-expression networks were built using the WGCNA R package (Langfelder and Horvath, 2008, 2012) using the normalized (TMM) and filtered expression data set (as previously described). Networks were built using the “signed” networkType parameter, enabling to capture the direction of the expression variation and grouping genes with the same direction variation in gene expression. This parameter is advised to identify biologically meaningful modules (van Dam et al., 2018). According to the mean connectivity and the scale free topology index curves obtained, a power of 20 was used for this analysis. A mergeheight parameter of 0.25 and a minimum module size of 30 were also considered. The modules obtained from this analysis were then tested for their enrichment in genes presenting differential expression patterns between the wild and domesticated pools.

## 3 Results

### 3.1 Nucleotide diversity patterns

For the analysis of polymorphism patterns the whole transcriptome of the wild and domesticated pools were analysed separately (Table 1 and Supplementary Table S12). Although a higher number of genes were retrieved for domesticated sorghum, a higher total number of polymorphic sites was called in the wild pool, suggestive of higher diversity in the wild populations (Table 1). Domesticated accessions showed a higher fixation index than wild sorghum (*F*=0.99 and *F*=0.42 respectively), in agreement with a higher rate of self-fertilization generally observed in the crop (e.g. Djè et al., 2004; Muraya et al., 2011).

The principal component analysis retrieved two genetic groups corresponding well to the wild and the domesticated pools (Figure 1). The only exception was the domesticated accession EC9 (SSM1057) that clustered with the wild samples. This accession belongs to the race *guinea margaritiferum*, which is cultivated but genetically close to the wild pool (Sagnard et al., 2011; Billot et al., 2013; Mace et al., 2013).

**Figure 1.**
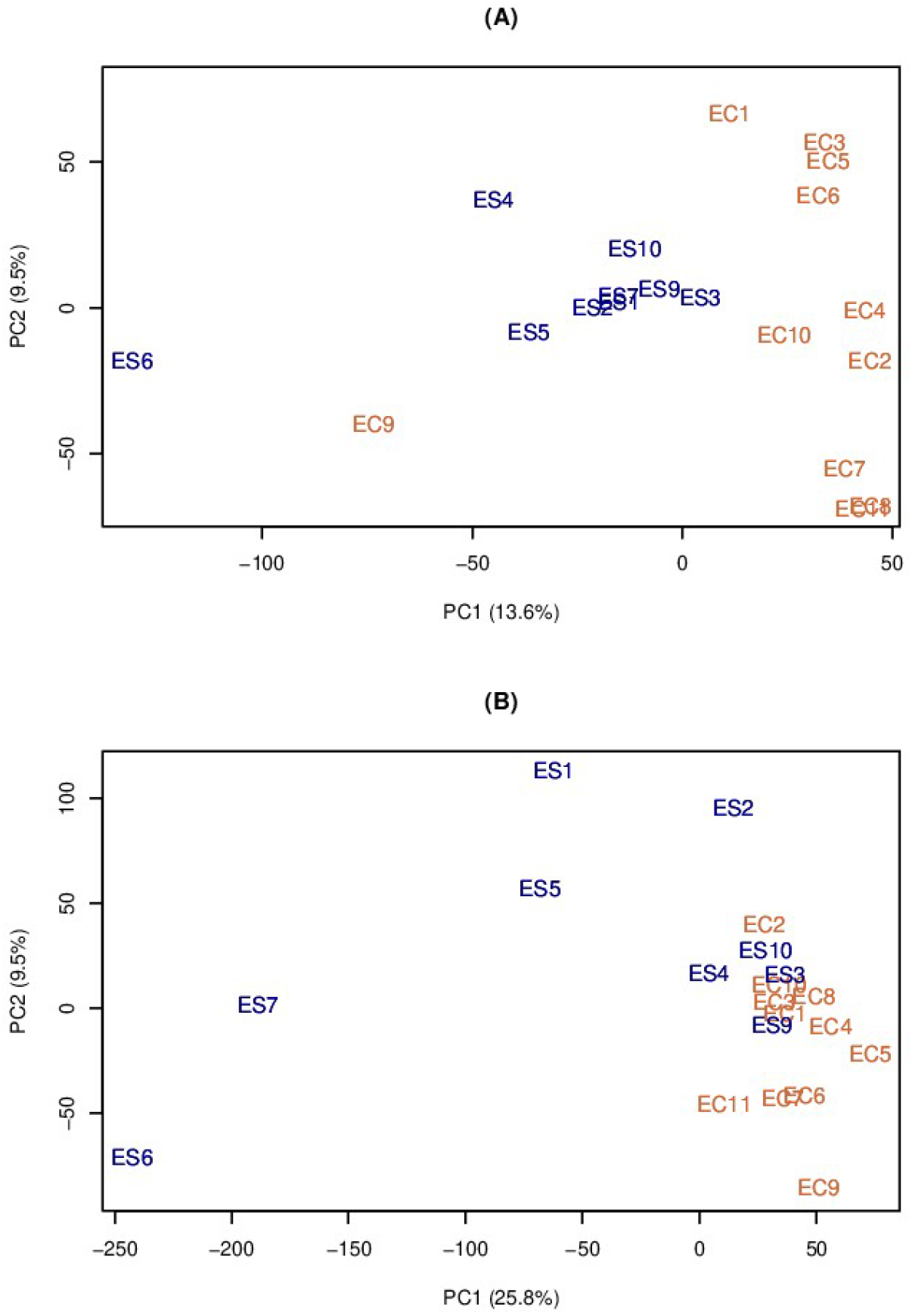
Principal Component analysis of polymorphism (A) and expression data (B) for wild (blue) and domesticated (orange) accessions of *Sorghum bicolor*. ES: wild; EC: domesticate.

Domesticated sorghum harbored 20-30% less nucleotide diversity than the wild counterpart, in terms of number of polymorphic sites and nucleotide diversity (Kolmogorov-Smirnov test p < 2.2. e-16; Table 1). Non-synonymous polymorphism (*π*_N_) was six times lower than synonymous polymorphism (*π*_S_) in both domesticated and wild pools, in agreement with previous estimates for domesticated sorghum from Hamblin *et al*. (2006). We did not find evidence of accumulation of slightly deleterious mutations in domesticated sorghum at the genome scale, since the ratio of non-synonymous to synonymous polymorphism (*π*_N_/*π*_S_) across genes was similar in the wild and domesticated pools (mean values 0.16 and 0.15, respectively, Table 1).

We also estimated Tajima’s *D*, which measures the departure of nucleotide diversity from neutrality due to selective or demographic processes. Domesticated and wild sorghum showed comparable close-to-zero mean Tajima’s D values across genes (Table 1). Similar values were found genomewide (Mace et al., 2013). The domesticated pool showed higher variance of Tajima’s D gene estimates (Table 1). This can point to a higher number of genes departing from neutral equilibrium, although a higher stochastic variance associated with lower polymorphism cannot be ruled out.

We calculated the ratio *π*_crop_/*π*_wild_ on 7853 genes. We considered the mean value *π*_crop_/*π*_wild_ = 0.65 (Table 1) to correspond to the global loss of genetic diversity due to the domestication bottleneck. Taking this value as reference, single gene *π*_crop_/*π*_wild_ ratio significantly different from the domestication bottleneck expectation may point to genes whose diversity has changed because of selection. Since strong positive selection is expected to reduce diversity in the genomic region linked to the causal allele, *π*_crop_/*π*_wild_ < 0.65 would characterize genes under positive selection in the domesticate. In particular, strong domestication candidates are genes that are monomorphic in the domesticate and polymorphic in the wild pool. Conversely, *π*_crop_/*π*_wild_ > 0.65 may indicate a diversity increase following domestication, either due to other selective dynamics (e.g. balancing selection or diversifying selection across varieties) or to a relaxation of selection in the domesticated pool compared to the wild one. We found that 1734 genes (22%) showed *π*_crop_/*π*_wild_ = 0 and 2500 (32%) had *π*_crop_/*π*_wild_ > 1.

Finally, we estimated the genetic differentiation between domesticated and wild sorghum by calculating *F*_ST_ on a gene by gene basis. Overall differentiation was quite low (mean *F*_ST_=0.07, Table 1), and did not change when genes with less than 5 polymorphic sites were excluded. Our estimate is lower than previous estimates (*F*_ST_=0.13 on SSRs, Casa *et al*. 2005). Two reasons can be proposed here to explain this discrepancy. Firstly, the coding regions analyzed in this study could have experienced a lower neutral drift than the microsatellite markers used by Casa *et al*. (2005). Secondly, it can emerge from the different compositions of the panels used. Indeed, we included a *guinea margaritiferum* accession in our domesticated pool whereas such type of accession was, to the best of our knowledge, not used in the analysis of Casa *et al*. (2005).

### 3.2 Expression diversity patterns

Reads counts were found for 38,885 genes, with 95% of them with read counts in either the domesticated or the wild pool. The analysis of gene expression levels was done on 24,646 genes meeting the criterion of at least 1 read count per million in at least 5 samples (Supplementary Table S12). Overall, our sampling strategy (3 different tissues) allowed accessing more than 70 % of the sorghum transcriptome. Similarly to the nucleotide diversity information, to compare the diversity in expression between domesticated and wild sorghum, we performed a principal component analysis and calculated the coefficient of variation (CV) on normalized read counts. PC1 separated the domesticated and wild pools, explaining 25.8% of the read counts diversity (Figure 1). The domesticated pool harbored 84% of the wild pool expression variability (mean CV_crop_=0.63 [95% quantiles 0.13; 1.98], mean CV_wild_=0.75 [95% quantiles 0.16; 2.04]), which suggests that domestication in sorghum involved also a loss of expression diversity, besides the loss of polymorphism reported above. We found that the CV reduction is higher (at least 25%) for candidate genes harboring *F*_ST_ excess and diversity reduction (potential signatures of selection) at the nucleotide level irrespective of the statistic used to identify them. This signal is stronger for genes with extremely high domesticate-wild differentiation. The CV loss was 34%, 32% and 29% in the 99%, 95% and 90% percentile of *F*_ST_ values respectively (Supplementary Figure S1). To exclude that this stronger CV reduction was simply due to a lower nucleotide diversity in *F*_ST_ outlier genes by comparison with non-outlier ones, we calculated the difference CV_wild_-CV_crop_ for 1000 random samples of non-outlier genes with equal or lower diversity level (*π*) than outlier genes. Wild to domesticate CV reduction in *F*_ST_. outliers was significantly higher than the CV reduction obtained with the random samples for the three *F*_ST_ percentiles (p-value < 0.001 for 95% and 90% and p-value < 0.01 for 99%; Figure 2).

**Figure 2.**
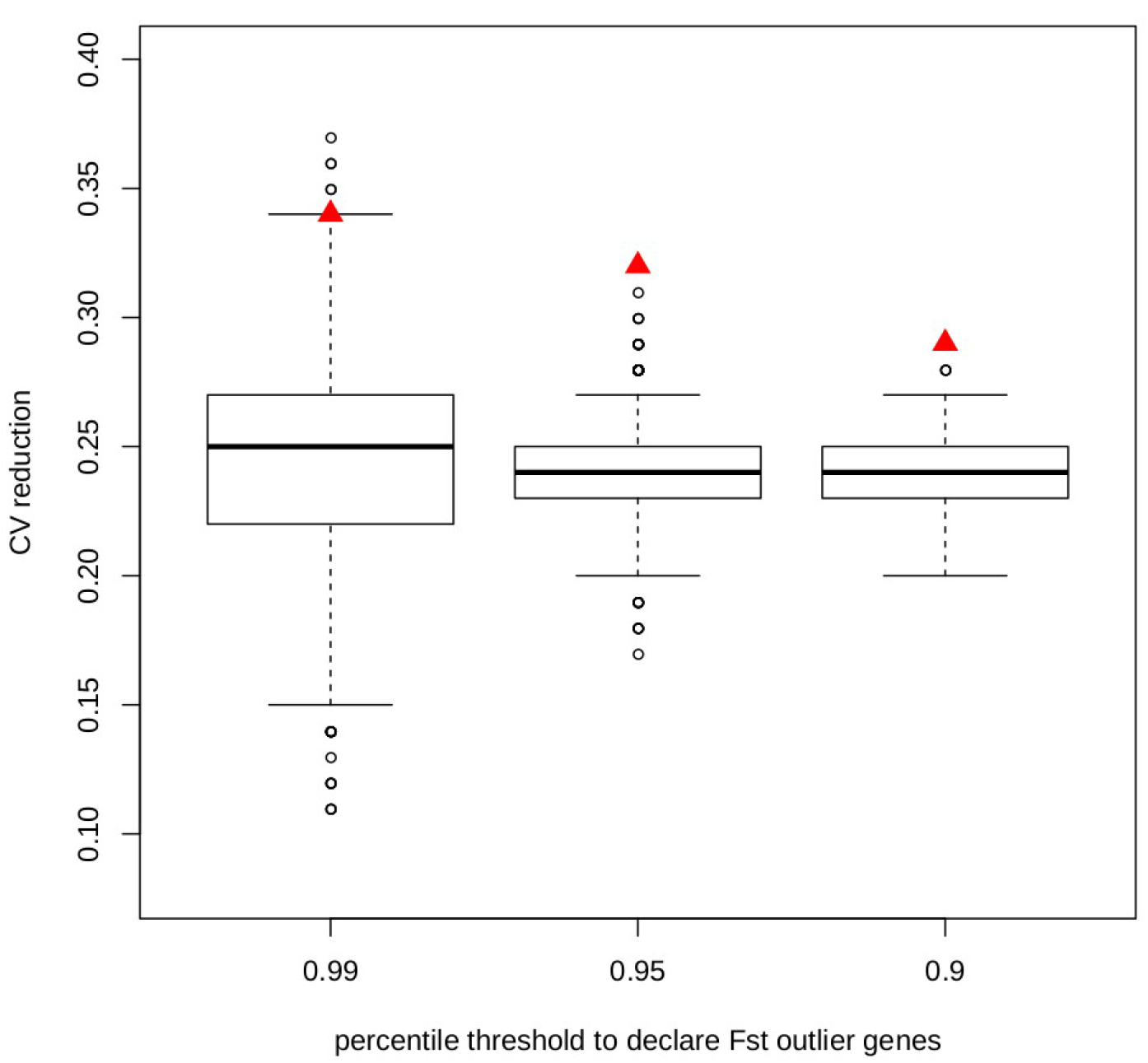
Observed (red triangles) and expected (black boxplots) wild to domesticate reduction of the coefficient of variation in expression (CV). Observed values were calculated on genes with extreme values of domesticate-wild differentiation (*F*_ST_ outliers) at different *F*_ST_. thresholds: 99%, 95% and 90% upper percentiles. Expected values were calculated on 1000 random samples of non-outliers genes with sample size equal to the number of outliers at each *F*_ST_. threshold and equal or lower levels of diversity *π*. Sample sizes: 92, 457 and 913 respectively. CV reduction was calculated as 1-(mean CV_CROP_ / mean CV_WILD_).

We tested for significantly different expression levels between the wild and domesticated pools, to identify genes whose change in expression may have been driven by the process of domestication. We found 949 differentially expressed (DE) genes at a FDR of 1% (Table S3; Supplementary Figure S2). The difference in expression levels corresponded to ļlogFCļ > 0.55 (logFC range: −9 to 14). Interestingly, among the DE genes, 82% were up-regulated in wild sorghum, while we did not observe any bias in the direction of expression change in the whole gene set (χ^2^ test, p < 2e-16). This result is qualitatively similar for a FDR of 5% (among 2291 DE genes, 72% were up-regulated in wild sorghum; Supplementary Figure S3).

### 3.3 Comparison of expression and nucleotide diversity patterns

We first compared expression and nucleotide patterns across all genes. We found that gene polymorphism (either estimated with *π*_S_, *π*_N_ or *π*_N_/*π*_S_) was negatively correlated with expression (Table 2). Negative correlation between expression level and *π*_N_ or *π*_N_/*π*_S_ is expected if higher expressed genes experience stronger selection against new deleterious mutations, whereas this pattern for *π*s may suggest other processes in action (e.g. selection on code usage). For *π*_N_/*π*_S_, the correlation was less negative for the domesticated accessions, which may be the effect of weaker purifying selection and domestication load experienced in the domesticated pool. Indeed, the analysis of variance showed that the relationship between polymorphism and expression is significantly different between the two genetic pools for all *π* estimates (Table S2); a difference potentially due to the process of domestication.

**Table 2.** Correlations of polymorphism estimates and expression levels.

|  |  | **Slope** | **adjR<sup>2</sup>** | **p-value** |
| --- | --- | --- | --- | --- |
| $\pi_s$ | Domestic | -0.39 | 0.13 | <2e-16 |
|  | Wild | -0.26 | 0.06 | <2e-16 |
| $\pi_N$ | Domestic | -0.54 | 0.24 | <2e-16 |
|  | Wild | -0.44 | 0.18 | <2e-16 |
| $\pi_N/\pi_s$ | Domestic | -0.14 | 0.014 | <2e-12 |
|  | Wild | -0.19 | 0.03 | <2e-16 |

$\pi$ : nucleotide diversity at synonymous sites ( $\pi_s$ ) and non-synonymous sites ( $\pi_N$ )

Then we looked at specific features of DE vs non-DE genes to identify signatures associated with differential expression. DE genes showed values of *π*_N_ and *π*_N_/*π*_S_ lower than non-DE genes in both domesticated and wild pools (Table 3), suggesting stronger purifying selection on DE genes, although the distributions were significantly different only for *π*_N_ in domesticated sorghum, (Kolmogorov-Smirnov test p-value = 0.008). DE genes were also characterized by significantly lower values of domesticate to wild *π* ratio than non-DE genes (p-value = 0.02 and 0.04 for *π*_N_ and *π*_S_, respectively). This difference was largely driven by genes that are monomorphic in domesticated sorghum, which may represent genomic regions targeted by human-driven directional selection. DE genes also showed higher Tajima’s D values than non-DE genes, but the difference was significant only in the wild pool (p-value = 0.024). We did not find significant differences between DE and non-DE genes for *π*_S_ or crop-wild *F*_ST_ (Table 3). Enrichment tests were, in general, consistent with distribution comparisons. DE genes were significantly enriched of *π*_crop_/*π*_wild_ values lower than 0.1 (corresponding to the lowest 20th percentile; Fisher exact test p-value = 0.03) and *F*_ST_>=0.35 (top 5th percentile; p-value=0.0014). Furthermore, among DE genes, 187 showed *F*_ST_= 0; these genes are good candidates of domestication-driven expression divergence involving only regulation changes without coding sequence divergence.

**Table 3.** Comparison of polymorphism patterns between genes significantly differentially expressed (DE) at 1% FDR and non differentially expressed genes (non-DE) between wild and domesticated *Sorghum bicolor*.

|  | DE genes (n=949) |  | Non-DE genes (n=23697) |  |
| --- | --- | --- | --- | --- |
|  | Wild | Domesticate | Wild | Domesticate |
| **No. upregulated genes** | 773<sup>c</sup> | 176 | 11287 | 12410 |
| **mean $\pi_S$<sup>a</sup>**<br>**[95% CI]** | 4.67<br>[0.03; 20.5] | 3.73<br>[0.03; 24.9] | 4.09<br>[0.03; 22.7] | 3.28<br>[0.03; 21.5] |
| **mean $\pi_N$<sup>a</sup>**<br>**[95% CI]** | 0.61<br>[0.03; 4.1] | 0.48**<br>[0.03; 3.8] | 0.65<br>[0.03; 4.5] | 0.5**<br>[0.03; 3.4] |
| **mean $\pi_N/\pi_S$<sup>a</sup>**<br>**[95% CI]** | 0.13<br>[0; 1.1] | 0.13<br>[0; 1] | 0.16<br>[0; 1.1] | 0.15<br>[0; 0.97] |
| **Tajima's D<sup>b</sup>**<br>**Median [95% quantiles]** | -0.25*<br>[-1.5; 1.5] | -0.19<br>[-1.6; 1.7] | -0.31*<br>[-1.5; 1.4] | -0.26<br>[-1.7; 1.6] |
| **Median**<br>**$\pi_{N-CROP}/\pi_{N-WILD}$** | 0.799* |  | 0.801* |  |
| **Median**<br>**$\pi_{S-CROP}/\pi_{S-WILD}$** | 0.767* |  | 0.769* |  |
| **Mean $F_{ST}$<sup>b</sup>**<br>**[95% quantiles]** | 0.09 [0; 0.4] |  | 0.07 [0; 0.3] |  |

<sup>a</sup> calculated on 469 genes with polymorphism estimates
<sup>b</sup> calculated on 9867 genes with polymorphism estimates
<sup>c</sup> Among DE genes, the number of genes up-regulated in the wild pool is significantly higher than among non-DE genes ( $\chi^2$ test p-value < 0.001)
\* distribution significantly different in DE vs non-DE genes (Kolmogorov-Smirnov test \*p-value < 0.05, \*\* p-value < 0.01)
$\pi$ : nucleotide diversity ( $\times 10^{-3}$ ). synonymous sites ( $\pi_S$ ), non-synonymous sites ( $\pi_N$ )

### 3.4 Biological functions of differentially expressed genes

Gene ontology (GO) analysis of DE genes revealed that terms related to photosynthesis were largely overrepresented among genes with reduced expression in the domesticated relative to the wild sorghum (n=51 genes, Table S4 and Supplementary Figure S4). Different metabolic functions were instead enriched in DE genes upregulated in the domesticate, the more recurrent being related to cytoskeleton organization and intracellular membrane transport (Table S5 and Supplementary Figure S5). Genes upregulated in the domesticated pool were enriched also in GO terms associated with Auxin responsive factors (ARF, GO:0005086 and GO:0032012). Gene functional annotation for all genes analysed in this study is provided in Supplementary Table S11.

We tested DE enrichment in Sorghum transcription factors (Plant Transcription Factor Database http://planttfdb.cbi.pku.edu.cn/index.php?sp=Sbi) and genes from the Sorghum expression Atlas (Shakoor et al., 2014). We found that DE genes were significantly depleted of transcription factors (p-value=0.009) and of genes reported to be stably expressed across tissues (p-value= 3.462e-05) compared to non DE genes.

We also compared our DE gene list with the candidate genes involved in trait domestication and improvement of sorghum according to previous studies. We found 59 genes harboring either a domestication or improvement signature as reported in Mace *et al*. (2013) and Lai *et al*. (2018) (Table S6).

Several DE genes belonged to known metabolic pathways in sorghum. Among them there were two genes involved in the sucrose metabolism and 4 involved in the phenylpropanoyl monolignol pathway according to Shakoor *et al*. (2014), but these metabolisms were not more represented in DE than in non DE genes. Other DE genes were previously known to be involved in the biochemical pathways associated with grain development and filling phases (Table S6), such as starch synthesis (Sobic.001G100000, Campbell et al., 2016), polyphenols synthesis (4 genes, Rhodes et al., 2014) and more generally grain composition (9 genes, Rhodes et al., 2017).

### 3.5 Gene co-expression networks

Gene co-expression network analyses revealed 21 clusters of genes sharing similar expression patterns (Table S7). Four of these clusters were enriched in differentially expressed genes between the wild and domesticated pools. Three were characterized by up-regulation and higher variability in wild sorghum (yellow: 2051 genes, magenta: 173 genes and cyan: 92 genes) and one was characterized by up-regulation and higher variability in domesticated sorghum (greenyellow: 133 genes) (Table S8). Among these four clusters, two were also enriched in domestication and improvement candidate genes previously identified through whole genome sequencing of 44 sorghum accessions including 7 wild and weedy genotypes (Mace et al., 2013) (Table S9 and S10). More specifically, the yellow and greenyellow modules were enriched in domestication genes whereas the greenyellow module was also enriched in improvement genes. Interestingly, the greenyellow module that harbored an enrichment in improvement genes presented up-regulation in the domesticated pool compared to the wild pool. In addition, the black (271 genes) and the lightcyan (45 genes) modules harbored expression patterns revealing the specific properties of the *guinea margaritiferum* accession (SSM1057). In the black module SSM1057 harbored up-regulation in comparison to all the other accessions analyzed, whereas in the lightcyan module SSM1057 presented an expression pattern more closely related to the wild pool (although it is a domesticated sorghum).

## 4 Discussion

In this study, we have performed a comparison of nucleotide and expression patterns between wild and domesticated forms of sorghum, to contribute to the understanding of the genetic circumstances of wild-to-domesticate transition in this major cereal. Much of the genome is not expressed. By targeting the full transcriptome, we focused on genes actually expressed, narrowing the analysis of interest to the part of diversity potentially most relevant. We found different lines of evidence documenting the genetic changes likely associated with the domestication and improvement process in sorghum.

### 4.1 Down-regulation and diversity loss during sorghum domestication

Our genome-wide screen of expressed genes showed a loss of diversity in the domesticated pool up to 30%, both in nucleotide and in expression diversity, supporting the general model of sorghum domestication from a reduced representation of the wild pool (e.g. Hamblin et al., 2006) and/or serial founder effects during crop evolution (Wang et al., 2017; Allaby et al., 2019). Our estimate of nucleotide diversity loss was higher than previous estimates for domesticated sorghum based on nuclear microsatellite markers (e.g. 14%, Casa et al., 2005), but lower than the loss recorded in other main crops with selfing or mixed mating systems. For instance, it was estimated a 62% nucleotide diversity reduction in rice (Caicedo et al., 2007), 72% in common beans (Bitocchi et al., 2013) and 50% in soybean (Kim et al., 2020). Domesticated sorghum also showed lower expression variation than the wild accessions, a finding common to other crops such as maize (Lemmon et al., 2014) and common bean (Bellucci et al., 2014) but also animal domesticates (Liu et al., 2019). We demonstrated that the loss of expression diversity is not merely a consequence of the observed loss of coding nucleotide diversity. In fact, the genes with strongest signatures of selection showed greater loss of expression variation, which suggests that selection has driven strongest changes in particular genes, a finding common to other domesticate species (Bellucci et al., 2014; Liu et al., 2019).

However, in contrast with a previous work on sorghum (Smith et al., 2019), and evidence in other plant and animal domesticates (Liu et al., 2017; Moyers et al., 2018), we didn’t find clear evidence of accumulation of deleterious load in the domesticated pool. Interestingly, a recent study in soybean found a genome wide reduction of the deleterious alleles in domesticated soybean relative to wild soybean (Kim et al., 2020). These authors suggested that the increase of self crossing associated with domestication may have helped purging the deleterious load, partly by exposing to selection deleterious variants in homozygous state, a process suggested also in cassava (Ramu et al., 2017) and grape (Zhou et al., 2017). A similar process can be at play in sorghum, where the domesticated pool generally shows a higher selfing rate than the wild pool (as observed in this study with the F index and also in Djè et al., 2004; Muraya et al., 2011; Sagnard et al., 2011). Further, gene flow from wild relatives (and vice versa) is considered frequent across the domesticate distribution (Barnaud et al., 2009; Mutegi et al., 2011, 2012; Sagnard et al., 2011) and has probably strongly contributed to the genetic makeup of the crop pool during the domestication process. Gene flow from wild sorghum has most likely counterbalanced drift and divergence genome-wide in the domesticated pool.

In nature, changes of gene expression are responsible for a great part of phenotypic divergence not explained by amino acid divergence (Carroll, 2008). It is also likely that part of the phenotypic evolution associated with the domestication process has involved gene regulation without changes in the coding sequence. This appears to be the case in sorghum, where many DE genes do not show different nucleotide diversity patterns between the domesticated and the wild pools, suggesting that domestication left a footprint in expression but not in protein composition for these genes.

Our differential expression analysis indicated that domestication largely favored the down-regulation of expression of multiple genes in sorghum crop (82% of DE genes). In agreement with our findings in sorghum, common bean and eggplant landraces showed downregulation of expression when compared with wild relatives, although the analysis was limited to the leaf transcriptome in both studies (Bellucci et al., 2014; Page et al., 2019). In maize, Lemmon *et al*. (2014), analyzing three different tissues (leaf, ear, stem) found the opposite trend, with differentially expressed genes showing higher expression of the maize allele more often than the teosinte allele. In contrast, Sauvage *et al*. (2017) found a similar number of down- and up-regulated genes in domesticated tomatoes based on the analysis of vegetative and reproductive tissues (leaf, flower and fruits). The lack of consistent patterns across different crops suggest species-specific processes and mechanisms for the evolution of the domesticated phenotype.

### 4.2 Biological significance of Differentially Expressed genes and genes harboring putative signature of selection at the nucleotide level

In this study, we found 949 differentially expressed genes between the wild and domesticated pools taking advantage of leaves, flowers and maturing seeds of sorghum. Among the genes significantly down-regulated in domesticated sorghum, genes with function associated with photosynthesis were disproportionately represented, suggesting that photosynthesis may be a key metabolic pathway contributing to the sorghum domestication syndrome. A number of studies have investigated the effect of domestication on photosynthesis metabolism, suggesting a shift in plant functional strategies over the course of domestication. For instance, higher net photosynthetic rate has been found in modern varieties of wheat compared to ancestral ones (Roucou et al., 2018) and in cultivated cassava compared with wild forms (Pujol et al., 2008), pointing to selection for increased photosynthetic efficiency during domestication and improvement. Genes significantly downregulated in domesticated sorghum suggested also that antioxidative responses have been weakened by domestication. Consistently with this scenario, it has been shown that ascorbate, an antioxidant widely associated with photosynthetic functions and stress tolerance, was reduced by domestication in some crop species, likely as a tradeoff of selection for higher fruit size and yield (Gest et al., 2013).

Modifications of the photosynthetic metabolism may be also a consequence of changes in the selective pressure experienced by the crop under some stresses. For instance, photosynthetic activity in leaves is overall reduced under drought stress (Osakabe et al., 2014). Some authors have suggested that the transition from wild habitats with unpredictable and scarcer water availability to agricultural environments with more regular and abundant water supplies led to a relaxation of selection for water use efficiency while favouring the increase of CO2 uptake and evaporative cooling (Milla et al., 2013 and references therein). It can be argued that domestication has led to a relaxation of selection for photosynthesis efficiency under drought stress, because of higher water availability in agricultural ecosystems than in the wild. This scenario would be consistent with positive selection for relatively higher photosynthetic capacity in cultivars adapted to water deficit environments, such as the case of upland rice compared with lowland rice (Zhang et al., 2016).

We found that genes upregulated in domesticated sorghum were enriched in Auxin responsive factors, which are very likely drivers of domestication-related phenotype. Genes involved in the response to auxin stimulus contribute to abiotic stress response in sorghum (Wang et al., 2010) and more generally is part of the auxin signaling machinery determining the development of branches and flowers in major crops such as maize (Galli et al., 2015). Auxin-responsive genes have been found under selection in sorghum, maize, rice and domesticated emmer (Mace et al., 2013; Meyer and Purugganan, 2013; Avni et al., 2017).

Altogether, our scan for selection signatures and differential expression identified several individual candidate genes particularly interesting for their potential role in the domestication and diversification of cultivated sorghum. Some of these genes are already known to contribute to important phenotypic traits in sorghum or have been previously identified as sorghum domestication genes. For instance, the gene Sobic.006G067700 (dwarf2) has been reported to control the internode length in domesticated sorghum and to have pleiotropic effects on panicle length, seed weight and leaf area (Hilley et al., 2017). This gene was upregulated in the domesticated pool (logFC=-0.95, FDR=4.14E-03), consistently with its involvement in the evolution of the domesticated phenotype. Another interesting gene is Sobic.004G272100, which encodes a phosphoribulokinase, an enzyme that catalyzes a key step in carbon fixation as part of the Calvin cycle (dark phase of photosynthesis). This gene has been found to be under selection in sorghum and maize (Lai et al., 2018). We found it was down-regulated in domesticated sorghum (logFC=1.86; FDR=0.0096), suggesting that human selection has targeted the reduction in expression level for this gene. In addition to the genes previously detected in the literature with evidence of signature of selection, our studies also provided a large list of additional genes that contributed or have been impacted by the domestication and improvement process. To further explore the list of genes that potentially contributed to these two key steps of crop evolution we studied their co-expression patterns.

### 4.3 Domestication and improvement steps involve major rewirings of gene expression

Gene co-expression network analyses allowed the identification of 4 clusters of genes (totalizing 2449 genes) significantly enriched in differentially expressed genes between wild and domesticated sorghum. Two of these clusters were also enriched in domestication and improvement genes previously identified in sorghum according to their nucleotide diversity patterns. These results reinforce the conclusion that can be drawn from the differential expression analysis indicating that domestication and improvement do not only affect the behavior of a few “domestication or improvement” genes but led to large rewirings of the transcriptome during the domestication event and the improvement process, as also underlined in tomato (Sauvage et al., 2017) and cotton (Gallagher et al., 2020). Whereas the “gene based” differential expression analyses can be limited by the sequencing depth achieved for each specific gene to detect significant expression differences, the co-expression gene network approach provides a complementary way to identify additional potentially relevant genes.

In the evolutionary history of sorghum, the *guinea margaritiferum* emergence remains until now quite unclear. Recent works proposed that they can result from a second and more recent domestication in West Africa (Sagnard et al., 2011; Mace et al., 2013). Our results indicated that the *guinea margaritiferum* accession presented a high similarity with the wild pool at the nucleotide level as previously reported, e.g. by Mace *et al*. (2013) and Morris *et al*. (2013). However, at the expression level, it behaved essentially as a domesticated genotype. Assuming a relatively recent domestication of the *guinea margaritiferum* sub-race, such results would indicate that the domestication process induced a rapid rewiring of the gene co-expression network and a much slower response at the genome-wide nucleotide level. It is also interesting to note that among the two clusters of genes for which an atypical behavior of the *guinea margaritiferum* was observed in comparison to the other domesticated genotypes (the black and lightcyan modules), there is the gene Sobic.001G341700, which has already been shown to contribute to the grain size in sorghum (Tao et al., 2017, 2020). This gene is an ortholog of Os DEP1 which has been proved to impact grain yield in rice (Huang et al., 2009).

## 5 Conclusion

This work highlighted the major impact of domestication not only on the nucleotide diversity but also on the transcriptome landscape. Indeed, it revealed a major rewiring of the gene co-expression networks between the wild and the crop pools. In addition to these global observations, these analyses also identified specific metabolic pathways (photosynthesis, auxin related mechanisms) that are likely to have contributed to the transition from the wild to the crop pools in sorghum. Finally, our results also contributed to a better understanding of the specific attributes of the *guinea margaritiferum* genotypes within the sorghum genetic diversity highlighting the specific role of the transcriptome regulation in the emergence of the “domesticated” ideotype.

## Acknowledgments

We wish to thank Pierre Roumet (INRAE, Montpellier, France) for useful inputs on the role of photosynthesis in domestication, and Benoit Nabholz (Montpellier University) for sharing scripts for sequence filtering. A first draft of this manuscript has been deposited as preprint, doi: https://doi.org/10.1101/2020.11.17.386268.

## Author Contributions

CB: Performed the analyses, interpreted the results, wrote the initial version of the manuscript, aggregated the suggestions of the co-authors and finalized the manuscript

AB: Managed the production of the tissues, took care of the RNA extraction and took part to the gene expression analyses

SG: Wrote and led the ARCAD project together with JD, contributed to the SNP calling section and contributed to the improvement of the manuscript

JD: Wrote and led the ARCAD project together with SG and contributed to the improvement of the manuscript

NT: Contributed to the interpretation of the differential expression results, contributed to the improvement of the manuscript

MD: Contributed to the interpretation of the nucleotide diversity results, and took part to the revision of the manuscript

DP: Contributed to the writing of the sorghum section of the ARCAD project and led the sorghum section of the BFF project. Selected the genotypes that have been used and took part in the analyses and writing of the manuscript.

## Supplementary Material

**Table S1**. Origin of Sorghum accessions used in this study

**Table S2**. Analysis of variance, with gene polymorphism explained by expression levels, gene pool and the interaction between them.

**Table S3**. Annotation, expression and nucleotide statistics for genes differentially expressed between domesticated and wild sorghum at 0.01 FDR

**Table S4**. Gene Ontology term enrichment in genes that were significantly up-regulated in wild sorghum at FDR < 0.01

**Table S5**. Gene Ontology term enrichment in genes up-regulated in domesticated sorghum at FDR < 0.01

**Table S6**. Differentially expressed genes found in common with other studies

**Table S7**: Membership of the 24646 genes to the 21 modules identified through the WGCNA analysis

**Table S8**: Enrichment of the modules detected through the WGCNA analysis in genes differentially expressed between the wild and domesticated pools

**Table S9**: Enrichment of the modules detected through the WGCNA analysis in genes detected by Mace et al., 2013 as harbouring signature of selection compatible with a domestication event

**Table S10**: Enrichment of the modules detected through the WGCNA analysis in genes detected by Mace et al., 2013 as harbouring signature of selection compatible with an improvement event.

**Table S11:** Functional annotation for all the 24646 genes analysed in this study.

**Table S12**: Expression and nucleotide statistics for all the 24646 genes analysed in this study.

**Figure S1.** Density distribution of the coefficient of variation (CV) in expression for two gene categories. On the left, genes with extreme values of crop-wild differentiation (*F*_ST_ outliers) identified at three percentile thresholds: 99%, 95%, 90% (from upper to bottom). On the right, all remnant genes (non-outliers). CV reduction was calculated as 1-(mean CV_CROP_ / mean CV_WILD_).

**Figure S2.** Plot of log-Fold Change (i.e. the log of the ratio of expression levels for each gene between wild and domesticated sorghum) against the log-concentration for count data (i.e. the overall average expression level for each gene across the two groups) for all genes analysed in this study (n= 24646), created with edgeR. Red: genes differentially expressed at 1% FDR (n=949).

**Figure S3**. Distribution of fold change (log scale) between wild and domesticated sorghum in different gene sets: all genes (n= 24646, top panel), genes differentially expressed at 5% FDR (n= 2291; middle) and genes differentially expressed at 1% FDR (n=949; bottom).

**Figure S4**. Graphical representation of GO terms associated with genes significantly upregulated in wild sorghum at 1% FDR (n=773) made with REVIGO.

**Figure S5**. Graphical representation of GO terms associated with genes significantly upregulated in domesticated sorghum at 1% FDR (n=176) made with REVIGO.

**Supplementary Text.** Script to perform the New Tuxedo pipeline, to calculate reads and transcript count from RNA seq data in Sorghum bicolor wild and domesticated accessions

