## Supplementary material for "The road to sorghum domestication: evidence from nucleotide diversity and gene expression patterns"

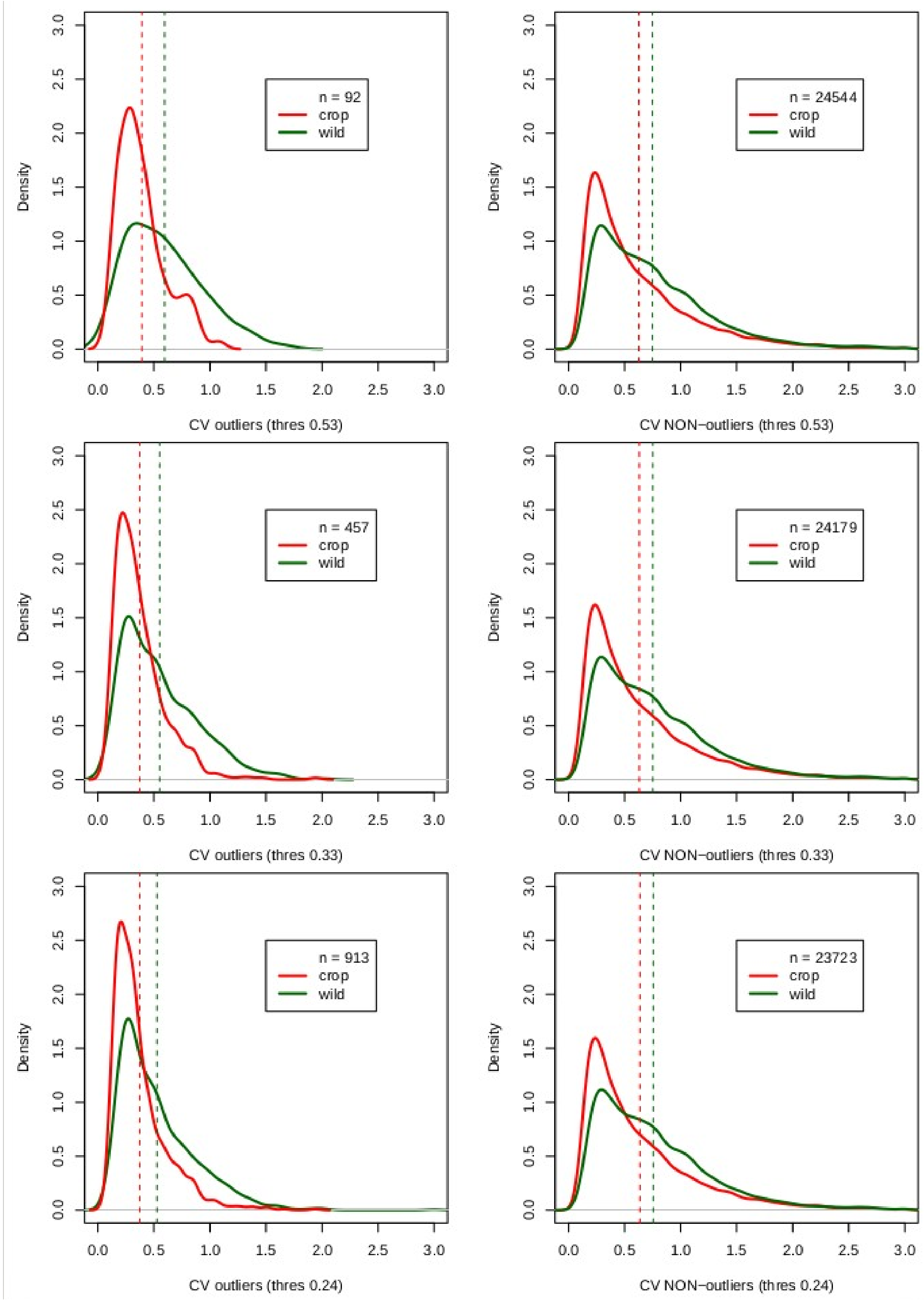


**Figure S1.** Density distribution of the coefficient of variation (CV) in expression for two gene categories. On the left, genes with extreme values of crop-wild differentiation (*F*ST outliers) identified at three percentile thresholds: 99%, 95%, 90% (from upper to bottom). On the right, all remnant genes (non-outliers). CV reduction was calculated as 1-(mean CVCROP / mean CVWILD).

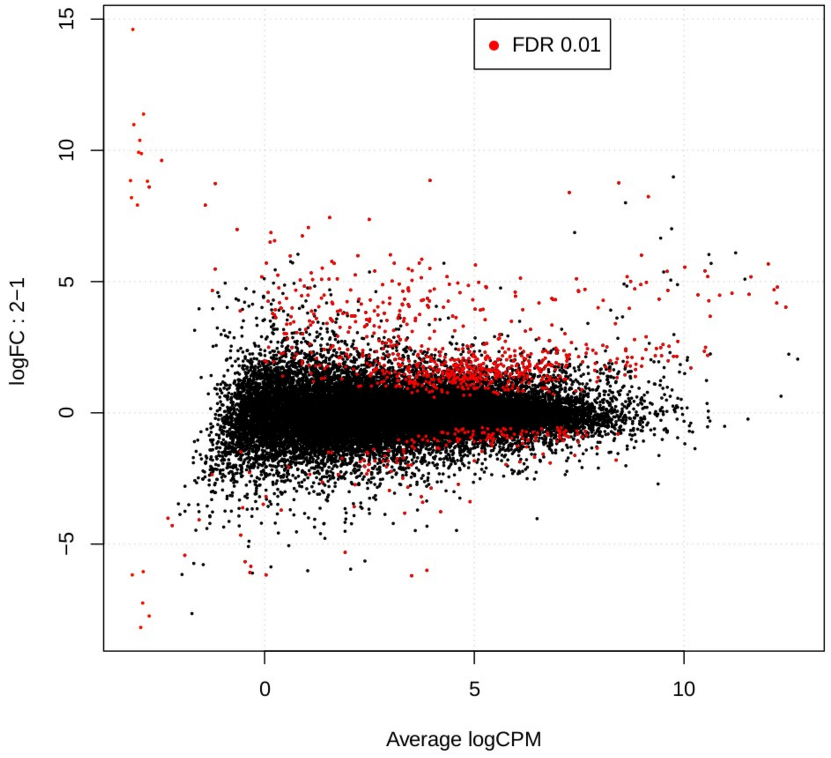


**Figure S2.** Plot of log-Fold Change (i.e. the log of the ratio of expression levels for each gene between wild and domesticated sorghum) against the log-concentration for count data (i.e. the overall average expression level for each gene across the two groups) for all genes analysed in this study (n= 24646), created with edgeR. Red: genes differentially expressed at 1% FDR (n=949).

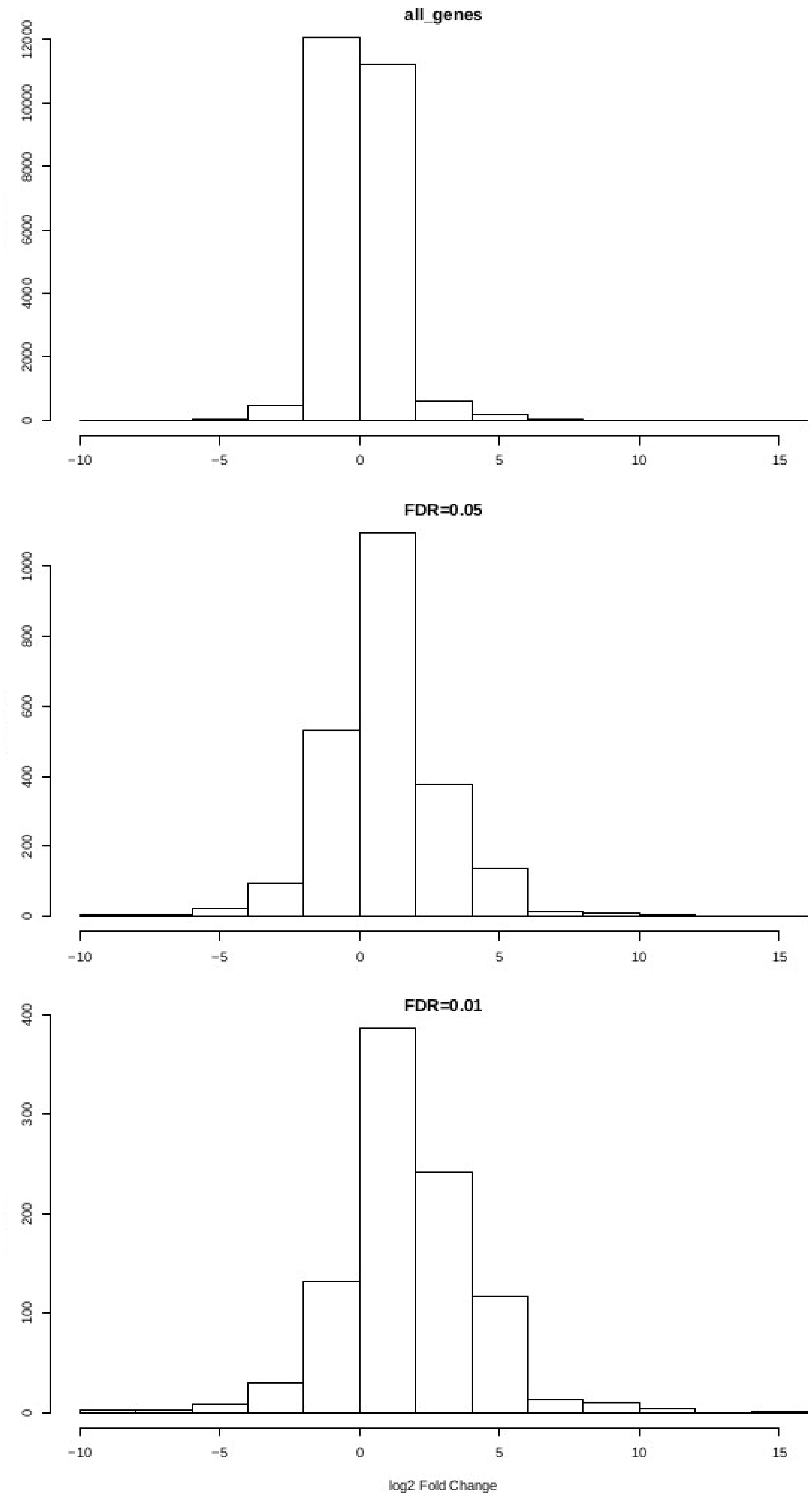


**Figure S3**. Distribution of fold change (log scale) between wild and domesticated sorghum in different gene sets: all genes (n= 24646, top panel), genes differentially expressed at 5% FDR (n= 2291; middle) and genes differentially expressed at 1% FDR (n=949; bottom).

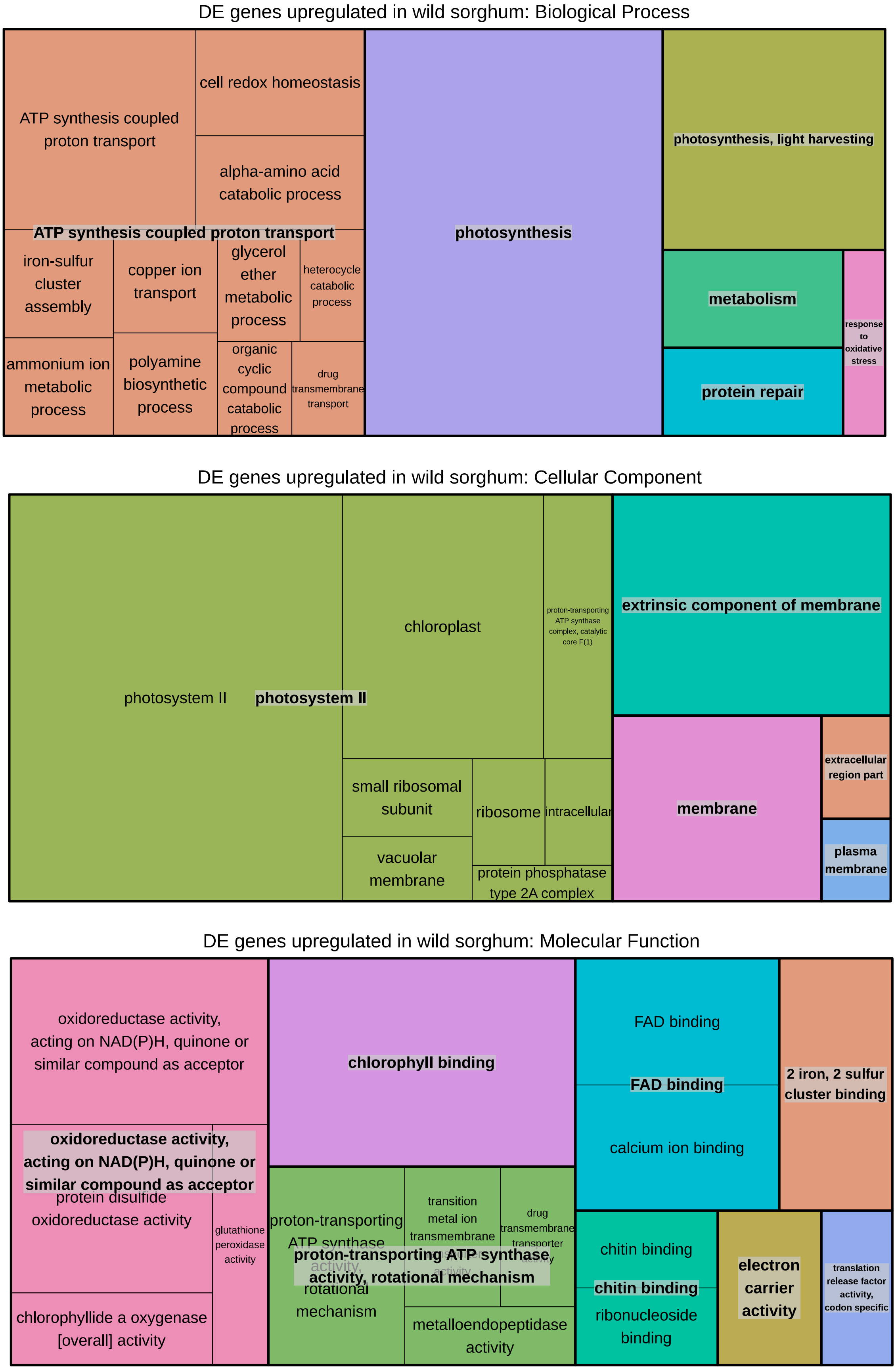


**Figure S4**. Graphical representation of GO terms associated with genes significantly upregulated in wild sorghum at 1% FDR (n=773) made with REVIGO.

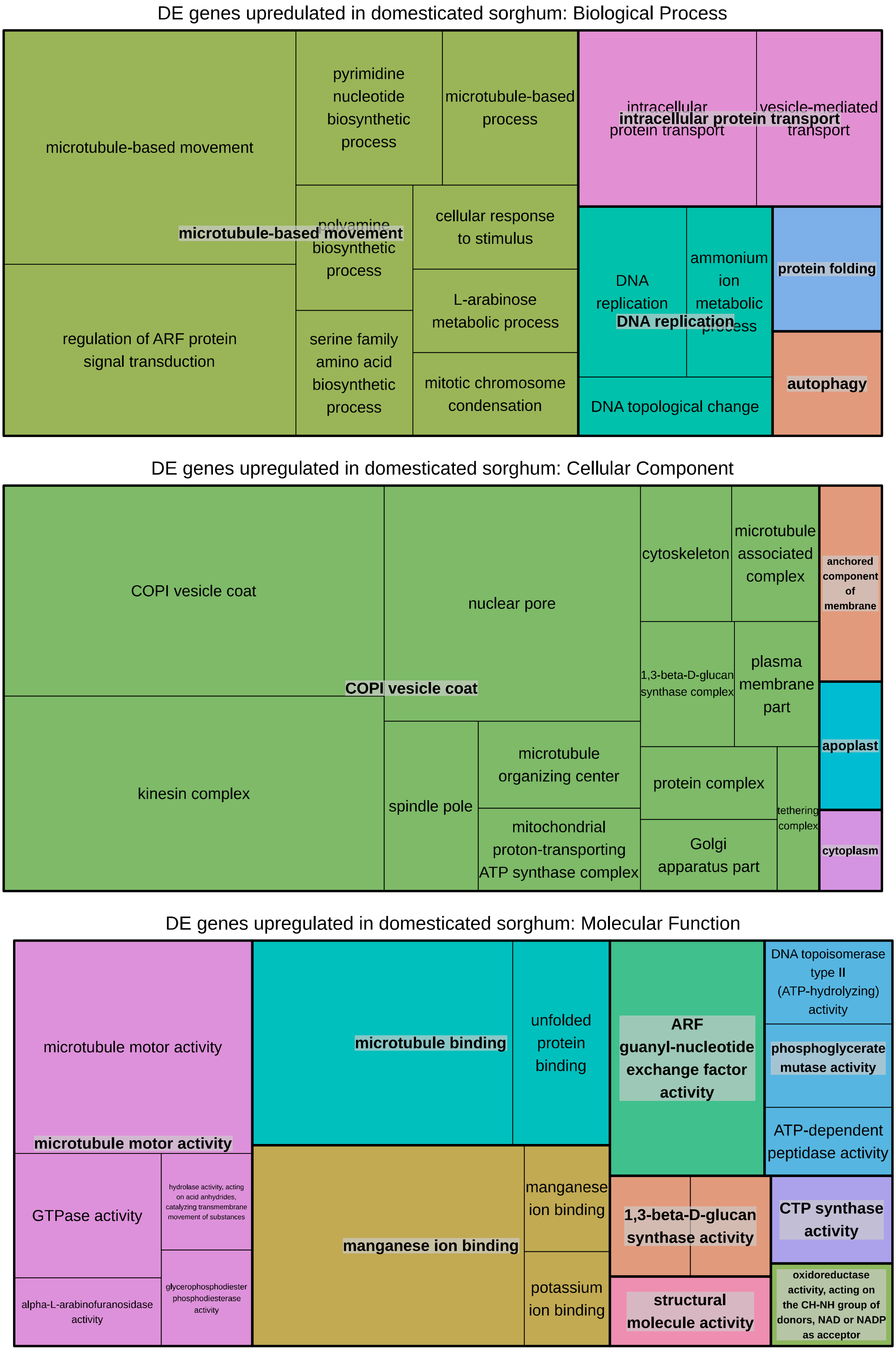


**Figure S5**. Graphical representation of GO terms associated with genes significantly upregulated in domesticated sorghum at 1% FDR (n=176) made with REVIGO.

**Supplementary Text 1**

Script to perform the New Tuxedo pipeline, to calculate reads and transcript count from RNA seq data in Sorghum bicolor wild and domesticated accessions

```
#!/bin/bash

##########################################################################################
#### Concetta Burgarella - Cirad
####
#### October 2017
####
#### New-tuxedo pipeline: pipeline to prepare gene and transcript count tables to perform a differential expression analysis
####
#### Sources: Pertea et al. 2016 Nature Protocols 11:1250, Angélique Berger (Cirad)
####
#### Use:
####
#### Put in the folder where analysis will be done: the scripts (diff_expression_pipeline.sh, geneIdtogeneName.pl, prepDE.py) and the sample.list file
####
#### Run the pipeline from the folder the script is with the following command line:
#### qsub -cwd -q normal.q -pe parallel_smp 20 -b y -M -m bea -V -N DE_pipeline.log ./diff_expression_pipeline.sh
####
##########################################################################################
# Prepare the sample.list in an automatic way for Arcad samples and BFF samples (for different samples adapt the command line or prepare the list manually)

# (run only once)
# cd /gs7k1/projects/arcad_data/sp1_final/sorghum/cleaned_data/
# ls *.gz >  /work/burgarella/file.list
# for file in $(cat /work/burgarella/file.list); do begin=$(echo `expr index "$file" E`); echo ${file:(($begin-1)):4}; done | tr -d '_' | tr -d '.' | sort > /work/burgarella/samples.list

##########################################################################################
# Load necessary modules

module load bioinfo/hisat2/2.0.5            # mapping utility
module load bioinfo/samtools                # sort and transform sam to bam
module load bioinfo/stringtie/1.3.3b        # assembling and quantifying genes and transcripts

##########################################################################################
# Set variables provided by the user

sample_list="samples.list";                 # list of sample names
echo "File with sample names: " $sample_list;
dos2unix "$samplelist" ;                    # check format compatibility

input_folder="/gs7k1/projects/arcad_data/sp1_final/sorghum/cleaned_data/";
 # localisation (folder) of qstat files
echo "Folder with qstat input files: " $input_folder;

ref_genome="/gs7k1/projects/BFF/sorghum_bicolor/Sorghum_genome_versions/annotation/formated_files/Sbicolor_313_v3.1.assembly.fna" ;  # fasta file of the reference genome for mapping (why not CDS only???)

ref_annotation="/gs7k1/projects/BFF/sorghum_bicolor/RNAseq/BFF/Sbicolor.gtf" ;
# annotation of the reference genome (it is a gff modified == .gtf prepared by A.B.)
```

```
gene_name="ARCAD_BFF" ;
# Name to be given to genes and transcripts not found in the genome annotation.

echo "Name to use for genes and transcripts:" $gene_name;

##############################################################################################
# Steps 1, 2, 3 (Pertea et al 2017) [performed in each sample folder (loop on samples)]

# for each sample of the list
for sample in $(cat "$sample_list") ; do
	echo "Sample treated now: " $sample ;
	echo "Input files: " "$input_folder"*"$sample"[._].* ;

	# Create a folder with the sample name
	mkdir "$sample" ;
	# change directory to the sample folder
	cd "$sample"	;
	# find and copy compressed files into the sample directory
	find "$input_folder" -iregex ".*$sample[._].*" -exec scp {} . \;
	# decompress tar.gz files
	tar -zxvf *gz	;

	# 1. Perform mapping
	echo "1. Mapping";
	hisat2 -p 20 -x "$ref_genome" -1 *forward* -2 *reverse* -U *single* -S "$sample".sam ;			#
SAM includes reads that failed to align
	echo "Mapping file created: " *.sam ;

	# 2. Sort .sam and transform to .bam (Sort alignments by leftmost coordinates, or by read name when -n is
used)
	echo "2. Sort and transform to bam";
	samtools sort -@ 8 -o "$sample".sorted.bam "$sample".sam ;   # -@ INT = Number of BAM compression
threads to use in addition to main thread [0]
	samtools flagstat "$sample".sorted.bam ;
	echo "Sorted bam file created: " *.sorted.bam ;

	# 3. assemble and quantify genes and transcripts (the output file of this step need a unique, ie per sample, file
name)
	echo "3. Assemble and quantify genes and transcripts";
	stringtie -p 10 -G "$ref_annotation" -o "$sample".gtf -l "$sample" "$sample".sorted.bam ;
# -G <ref_ann.gff> Use the reference annotation file (in GTF or GFF3 format) to guide the assembly process.
# The output will include expressed reference transcripts as well as any novel transcripts that are assembled.
# [-l <label> name prefix for output transcripts (default: MSTRG)]
	echo "Assembled genes and transcripts in gtf file: " *.gtf

	cd .. ;    # return to the parent directory

done

##############################################################################################
# Step 4. Merge individual gtf files (Pertea et al 2017) [to be done in the parent folder]

# create a file with the list of gtf file paths for all the samples wanted to be merged
ls */*.gtf > mergelist.txt ;	# gtf files created for current samples
ls /gs7k1/projects/BFF/sorghum_bicolor/RNAseq/BFF/ARCAD/Stringtie/BFF_2013/**/*.gtf    >> mergelist.txt ;	#
BFF samples
ls /gs7k1/projects/BFF/sorghum_bicolor/RNAseq/BFF/ARCAD/Stringtie/BFF_2014/**/*L*.gtf   >> mergelist.txt ;	#
BFF samples (some .gtf not wanted)
```

```
ls /gs7k1/projects/BFF/sorghum_bicolor/RNAseq/BFF/ARCAD/Stringtie/BFF_2015/**/*L*.gtf    >> mergelist.txt ;    #
BFF samples (some .gtf not wanted)

echo "Nb of samples included in the merged list:"
wc -l  mergelist.txt  # nb of files (should be 782)

# merge (in this step, transcripts not associated to a gene model included in teh reference annotation file are given a unic
specific name)
#   -l <label>       name prefix for output transcripts (default: MSTRG)

echo "4. Merge individual gtf files" ;
stringtie --merge -p 20 -G "$ref_annotation" -o "$gene_name"_merged.gtf -l "$gene_name" mergelist.txt ;
echo "Merged gtf file created:" *merged.gtf ;

##########################################################################################
# Step 5. Compare transcripts with the reference annotation (Pertea et al 2017) [to be done in the parent folder]
# adds to each transcripts a class code and the name of the transcript from the reference annotation file

echo "5. Compare transcripts with the reference annotation " ;
/gs7k1/projects/BFF/sorghum_bicolor/RNAseq/BFF/gffcompare/gffcompare -r "$ref_annotation" -o merged
"$gene_name"_merged.gtf ;

# change the order of designation for 2 columns (what is called "gene_name" becomes "gene_id" and viceversa)
# this is necessary to get again a format similar to ARCAD_BFF_merged.gtf that merged.annotated had changed
perl geneIdtogeneName.pl merged.annotated.gtf ;            # the output is merged.annotated.gtf.cleanId.gtf

#wc -l merged.annotated.gtf.cleanId.gtf  # 972623 (a bit less than BFF_782_merged.gtf...)

##########################################################################################
# 6. Estimate transcript abundance and store the results in tables to be read under the R environment [to be done in the
parent folder]
# needs .bam files, sample.file and merged.annotated.gtf.cleanId.gtf

echo "6. Estimate transcript abundance and store the results in tables" ;
mkdir abundance ;

# estimate abundance on a per sample basis and in a new folder per sample
for sample in $(cat "$sample_list") ; do
	echo "Sample treated now: " $sample ;
	echo "input files: " "$sample"/"$sample"*sorted.bam ;

	mkdir abundance/"$sample" ;

	# estimate transcript abundance
	# [ -e only estimate the abundance of given reference transcripts (requires -G)]
	# [ -B enable output of Ballgown table files which will be created in the same directory as the output GTF
(requires -G, -o recommended)]
	# [-l <label> name prefix for output transcripts (default: MSTRG)]
	stringtie -e -B -p 10 -l "$sample" -G merged.annotated.gtf.cleanId.gtf -o
abundance/"$sample"/"$sample"_abundance.gtf "$sample"/"$sample"*sorted.bam ;

done

# create tables of gene and transcript abundance with the phyton script provided by stringtie
./prepDE.py -i abundance -g "$gene_name"_gene_count.csv -t "$gene_name"_transcript_count.csv -p E -s
"$gene_name" ;

echo "Count table files generated:" *.csv ;

echo "END OF THE ANALYSIS"
```
